# Genotype-specific roles of small extracellular vesicles in modulating metronidazole resistance in *Giardia lamblia*

**DOI:** 10.1101/2025.01.17.633589

**Authors:** Gabriel Luna Pizarro, Jerónimo Laiolo, Nehuén Salas, Rocío G. Patolsky, Luciano Díaz Pérez, Camilo Cotelo, Constanza Feliziani, Andrea Silvana Rópolo, María Carolina Touz

## Abstract

*Giardia lamblia*, a eukaryotic intestinal parasite, produces small extracellular vesicles (sEVs) as a conserved evolutionary mechanism. This study investigates the functional role of sEVs in transferring drug-resistance traits among parasites. sEVs derived from metronidazole (MTZ)-resistant clones are shown to modify the expression of enzymes involved in MTZ metabolism and the production of reactive oxygen species (ROS) in recipient wild-type parasites. These changes significantly alter the drug sensitivity of recipient parasites. The transfer efficiency and phenotypic impact vary depending on the genetic background of the isolates, highlighting a genotype-specific mechanism. Our findings reveal that sEVs act as mediators of phenotypic adaptation in *G. lamblia*, enhancing parasite survival under drug-induced stress. This study underscores the importance of sEVs in drug-resistance dynamics and provides a basis for exploring therapeutic interventions targeting EV-mediated resistance in giardiasis.

**Highlights In brief:**

- Small extracellular vesicles (sEVs) from *Giardia lamblia* mediate genotype-specific MTZ resistance transfer.
- Small extracellular vesicles from drug-resistant clones (RsEVs) alter enzyme expression and reactive oxygen species (ROS) production in recipient trophozoites.
- The genetic background of *G. lamblia* isolates influences the effectiveness of resistance transfer.
- Findings provide insights into resistance mechanisms and potential targets for new giardiasis therapies.

## INTRODUCTION

Giardiasis, caused by the protozoan *Giardia lamblia* (syn. *G. intestinalis* or *G. duodenalis*), is the most common non-viral and non-bacterial diarrheal illness worldwide, leading to significant health issues such as weight loss, malnutrition, growth delays in children, delayed puberty, impaired cognitive development, and even premature death ^1,2^. The primary treatments for giardiasis are metronidazole (MTZ) and tinidazole, which belongs to the 5-nitroimidazole (5-NI) family. However, persistent infections can occur due to reinfection, inadequate drug dosage, immunosuppression, drug-resistant strains, and sequestration of *Giardia* in the gallbladder or pancreatic duct. Current treatments for parasitic infections are systemic, require long-term medication, come with side effects, and are ineffective against resistant strains. MTZ resistance in *Giardia* poses a significant challenge in treating giardiasis, and resistance has been observed across different medications ^3,4^.

According to a widely accepted model, nitro compounds are activated by reduction, producing toxic intermediates that cause oxidative stress ^5,6^. MTZ is a prodrug, pharmacologically inactive in its original form, which requires metabolic activation within the body to become its active form. Understanding the causes of MTZ resistance involves exploring the activation of nitro compounds and the subsequent detoxification pathways within the parasite. While resistance to nitro compounds is commonly observed both *in vitro* and *in vivo*, fresh resistant patient isolates are challenging to maintain in axenic culture. Consequently, most studies rely on generating resistant model strains *in vitro* and comparing them to isogenic wild type strains ^7^. Understanding the causes of MTZ resistance involves exploring the activation of nitro compounds and the subsequent detoxification pathways within the parasite. In this sense, transcriptional changes and proteome analysis have highlighted significant differences in gene expression between susceptible and resistant genotype A strains ^8–10^. Our working hypothesis is that strain genotypes can influence these changes, complicating the identification of common and specific resistance patterns across studies.

The genus *Giardia* is classified into eight species or groups depending on the host, with *G. lamblia* being the only species that infect humans (genotypes A and B) ^11,12^. Notable distinctions in pathogenicity and infectivity exist between genotypes A and B. Genotype B infections in humans are associated with more pronounced small intestinal inflammation, including villous shortening/atrophy and lamina propria inflammation, compared to genotype A (reviewed in ^13,14^). Furthermore, genotype B isolates induce more significant alterations in the intestinal mucosa and a reduction in the enzymatic activity of the brush border than genotype A ^15^. Additionally, genotype B is more resistant to reactive oxygen species (ROS) and nitric oxide (NO)^8,16–18^.

Extracellular vesicles (EVs) are membrane-bound particles released into the extracellular environment that mediate communication and information transfer between cells and organisms across all domains of life ^19^. These vesicles transport molecular cargo, including cytosolic and membrane proteins, lipids, and RNA ^20–22^. EVs play significant roles in normal physiology and pathogen-host interactions, spreading antigens and infectious agents. Initially studied for their functions in immune surveillance, EVs also facilitate various modes of communication between parasites ^23–27^. So far, studies on small and large EVs showed that they are mainly engaged in cellular stress responses and contribute to the resistance against chemotherapeutic drugs in eukaryotic cells, also recently reported in *Leishmania* ^28,29^. *Giardia* trophozoites produce different populations of EVs under various environmental conditions or biological stimuli, including large extracellular vesicles (mainly enriched in microvesicles - MVs) and small extracellular vesicles (sEVs, exosome-like) or the complete secretome (EVs plus free secreted proteins) ^30–32^. Our research group has previously examined the formation and release of small extracellular vesicles, from now called sEVs, in *Giardia* trophozoites and compared the types and quantities of small RNAs (sRNAs) in sEVs from strains with different genotypes ^33,34^.

In this work, we explored whether small extracellular vesicles from MTZ-resistant clones (RsEVs) produced *in vitro* could be involved in transmitting MTZ resistance intra- and inter-genotypes A and B. This study provides, for the first time, evidence that RsEVs can induce changes in the expression of enzymes involved in MTZ metabolism and ROS production and that these changes rely on the RsEVs information and the genotype of the naive, drug-sensitive parasites. This incorporation facilitates the rapid emergence of drug-sensitive or drug-resistant subpopulations based on genotype background, enabling swift adaptation to environmental stressors and enhancing survival under varying conditions.

## RESULTS

### Successful *in vitro* induction of metronidazole resistance in trophozoites of genotypes A and B

Metronidazole (MTZ) has been associated with the most significant resistance because it is the first line of treatment. To produce MTZ-resistant clones, wild type trophozoites from the WB/1267 (genotype A) and GS/M (genotype B) isolates were grown in a sublethal concentration of MTZ. After 730 days of continuous subculture with sublethal increasing concentrations of MTZ, strains WB/1267_MTZ_r and GS/M_MTZ_r were obtained and adapted to grow in the presence of MTZ at concentrations up to 6.4 μM and 10.5 μM, respectively, which were used in resistance studies. The survival of *G. lamblia* trophozoites was evaluated *in vitro* by microscopic observation, and the 3-(4,5-dimethylthiazol-2-yl)-2,5-diphenyltetrazolium bromide (MTT) colorimetric assay to determine the half-maximal inhibitory concentrations (IC_50_). IC_50_ was used to determine the concentration of MTZ necessary to inhibit the growth or viability of trophozoites by 50%. The IC_50_s of WB/1267 and GS/M wild type isolates were around 30 μM (Figure 1A), while the values for clones WB/1267_MTZ_r and GS/M_MTZ_r were 80 and 209 µM, respectively (Figure 1B). We also calculated the resistance fold (RF = IC_50_ MTZ Resistant / IC_50_ sensitive). The RF showed a marked increase in resistance to the antiparasitic drugs studied, with RF WB/1267 MTZ = 2.7 and RF GS/M MTZ = 6.8, respectively. Except for samples from clinical cases, laboratory-derived MTZ-resistant trophozoites from genotype B were not reported until this study. Therefore, this is the first report indicating that both genotypes that infect humans can develop resistance *in vitro* when exposed to suboptimal drug concentrations.

**Figure 1:**
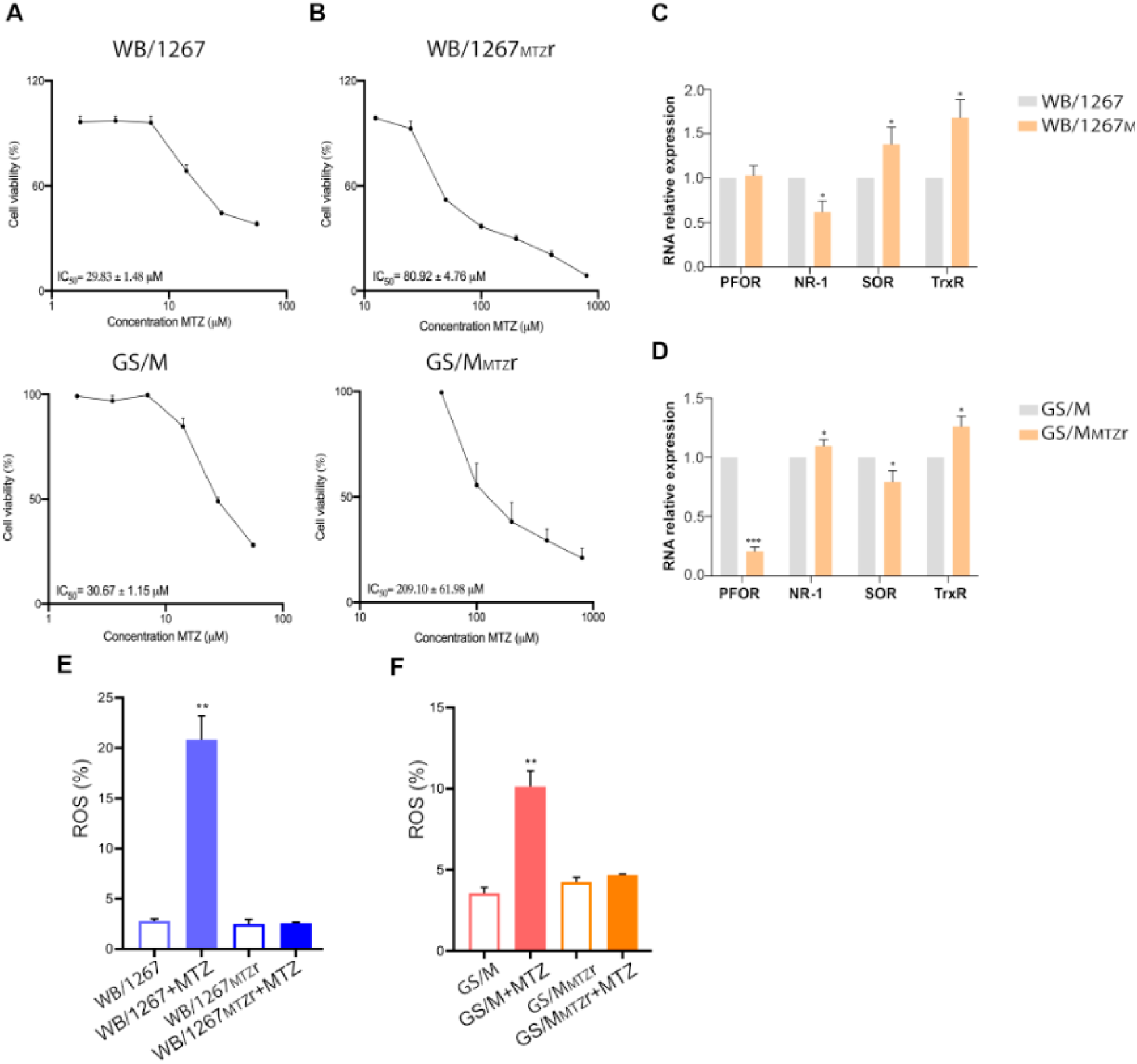
Evaluation of *Giardia lamblia* trophozoites’ induced metronidazole (MTZ) resistance mechanisms. (A-B) The IC_50_ values of MTZ for induced WB/1267_MTZ_r and GS/M_MTZ_r clones are significantly higher than the wild type isolates WB/1267 and GS/M isolates. (C-D) Relative mRNA expression levels of enzymes implicated in MTZ activation and detoxification, including pyruvate ferredoxin oxidoreductase (PFOR), nitroreductase-1 (NR-1), superoxide reductase (SOR), and thioredoxin reductase (TrxR), measured via qPCR. (E-F) Intracellular reactive oxygen species (ROS) production in response to 20 µM MTZ was assessed using H_2_DCFDA and flow cytometry and shown as a percentage compared with the control. *p < 0.05; **p < 0.01, ***p < 0.001.

### *G. lamblia* WB/1267MTZr and GS/MMTZr clones exhibit distinct patterns of enzyme expression involved in the MTZ’s metabolic pathways

MTZ resistance in *G. lamblia* involves the regulation of specific enzymes critical to the drug’s activation and detoxification processes. Due to MTZ’s shallow redox potential, its metabolization only quantitatively occurs in microaerophilic and anaerobic organisms with a strongly reductive physiology ^35^. Resistance often entails downregulating enzymes that convert MTZ to its toxic intermediates, such as pyruvate oxidoreductase (PFOR) and Nitroreductase-1 (NR-1), a ferredoxin-nitroreductase chimera as reviewed elsewhere ^8,36,37^. Conversely, resistance can be achieved by upregulating enzymes detoxifying MTZ or managing MTZ-induced damage, such as Nitroreductase-2 (NR-2) and superoxide reductase (SOR). SOR, commonly found in prokaryotic organisms, protects from oxidative stress by directly reducing MTZ to an inert amine ^38,39^. Reduced PFOR activity is associated with MTZ resistance, while increased SOR expression enhances detoxification. Thioredoxin reductase (TrxR) shows variable regulation across resistant strains, suggesting it plays a dual role in drug activation and oxidative stress management ^40^. To investigate the transcriptional levels of these critical enzymes involved in MTZ activation in WB/1267_MTZ_r and GS/M_MTZ_r clones, qPCR analysis of mRNA expression was performed for PFOR, NR-1, SOR, and TrxR, comparing these resistant lines to their susceptible wild type counterparts. The results for WB/1267_MTZ_r revealed that PFOR expression remained unchanged. In contrast, NR-1 expression was significantly reduced (Figure 1C). Additionally, SOR expression increased compared to WB/1267 wild type (Figure 1C). Analysis of TrxR expression showed a significant increase over the susceptible wild type line. These findings suggest that resistance in WB/1267_MTZ_r is due to the downregulation of NR-1, reducing MTZ cytotoxicity, and the upregulation of SOR, which helps render MTZ inert. The increased expression of TrxR indicates a protective role in these resistant clones. For the GS/M_MTZ_r clone, a remarkable decrease in PFOR mRNA expression and a significant increase in TrxR were observed. These findings may represent a combination of mechanisms involved in MTZ resistance in this clone (Figure 1D).

MTZ reduction forms highly reactive nitro radicals, damaging essential cellular components such as DNA, proteins, and lipids. Additionally, the nitro radicals react with available oxygen molecules, producing ROS. To analyze the intracellular generation of ROS in *Giardia* WB/1267_MTZ_r and GS/M_MTZ_r, 2′,7′-dichlorodihydrofluorescein diacetate (H_2_DCFDA), was used and measured by flow cytometry. The results showed that after adding 20 µM of MTZ, the WB/1267_MTZ_r and GS/M_MTZ_r clones produced significantly lower levels of ROS than their wild type isogenic isolates (Figure 1E-F). These results suggest that MTZ resistance in WB/1267_MTZ_r and GS/M_MTZ_r is linked to a reduced ability to generate ROS, typically produced during the drug’s activation process. These findings highlight the complex regulatory mechanisms underpinning MTZ resistance in *G. lamblia*, which reduce drug activation pathways and enhance detoxification and protective responses.

### The clones WB/1267_MTZ_r and GS/M_MTZ_r show MTZ-resistance stability after discontinuation of drug selection

Previous studies have shown that *in vitro*-generated MTZ-resistant *G. lamblia* genotype A trophozoites can revert to drug sensitivity after encystation or excystation and after several generations without selective pressure ^41^. However, increasing evidence suggests that clinically MTZ-resistant trophozoites can be transmitted between patients, indicating that resistance can be a stable, transmissible phenotype ^42,43^. To explore the potential for reversion to drug sensitivity and the durability of MTZ resistance in WB/1267_MTZ_r and GS/M_MTZ_r clones, we cultured these clones for 5, 10, or 15 days without drug selection. This timeline aligns with typical MTZ treatment regimens, lasting 5–10 days with one or two doses per day ^44^. MTZ IC_50_ values were calculated at each time point, showing that while resistance decreased over the 2 weeks without MTZ, the clones retained significantly higher resistance levels than their drug-sensitive parents and original isolates (Figure S1). These results suggest that while some reversion to drug sensitivity occurs without drug pressure, the parasites do not fully revert to their original sensitive state. This finding provides important insights into the long-term persistence of drug resistance in *Giardia*. It establishes a crucial basis for using these clones in downstream experiments, ensuring that observed effects result from experimental variables rather than variations in baseline resistance stability.

### MTZ-resistant WB/1267_MTZ_r and GS/M_MTZ_r clones release small extracellular vesicles (sEVs) that might act as carriers of drug resistance

Our recent study demonstrated that small extracellular vesicles (sEVs) from *G. lamblia* isolates WB and GS act as cell-to-cell shuttles for RNA ^33^. We hypothesized that RsEVs from metronidazole-resistant (MtzR) clones might transmit drug resistance by carrying molecules involved in resistance mechanisms. The sEV-enriched fractions were first characterized for size distribution and zeta potential (ZP) using nanoparticle tracking analysis (NTA) to evaluate our hypothesis. Figure 2A-B illustrates the mean particle size of sEVs derived from wild type and RsEVs from MTZ-resistant clones. The average sizes were similar across all samples, with mean diameters of 108.3 nm for WB/1267, 97.2 nm for GS/M, 114.6 nm for WB/1267_MTZ_r, and 107.0 nm for GS/M_MTZ_r (Figure 2C). The NTA analyses also revealed that the sEV preparations were free of contaminants and uniform in size. We also measured the zeta potential of these sEVs, which typically ranges from −10 mV to −50 mV for RsEVs (exosomal) and is influenced by surface charge, affecting particle aggregation tendencies ^45,46^. Measurement of the ZP of sEVs at 25°C in PBS showed mean values of −36.998 mV for WB/1267, −36.351 mV for GS/M, −38.457 mV for WB/1267_MTZ_r, and −38.834 mV for GS/M_MTZ_r (Figure 2C). These negative ZP values indicate that the sEVs exhibit electrostatic repulsion between particles, which helps prevent aggregation and supports colloidal stability. Transmission electron microscopy (TEM) was used to evaluate the size and morphology of the sEVs. The results showed that the sEVs were around 100 nm in size and had cup-shaped structures for all samples (Figure 2D), consistent with previous findings ^33^.

**Figure 2:**
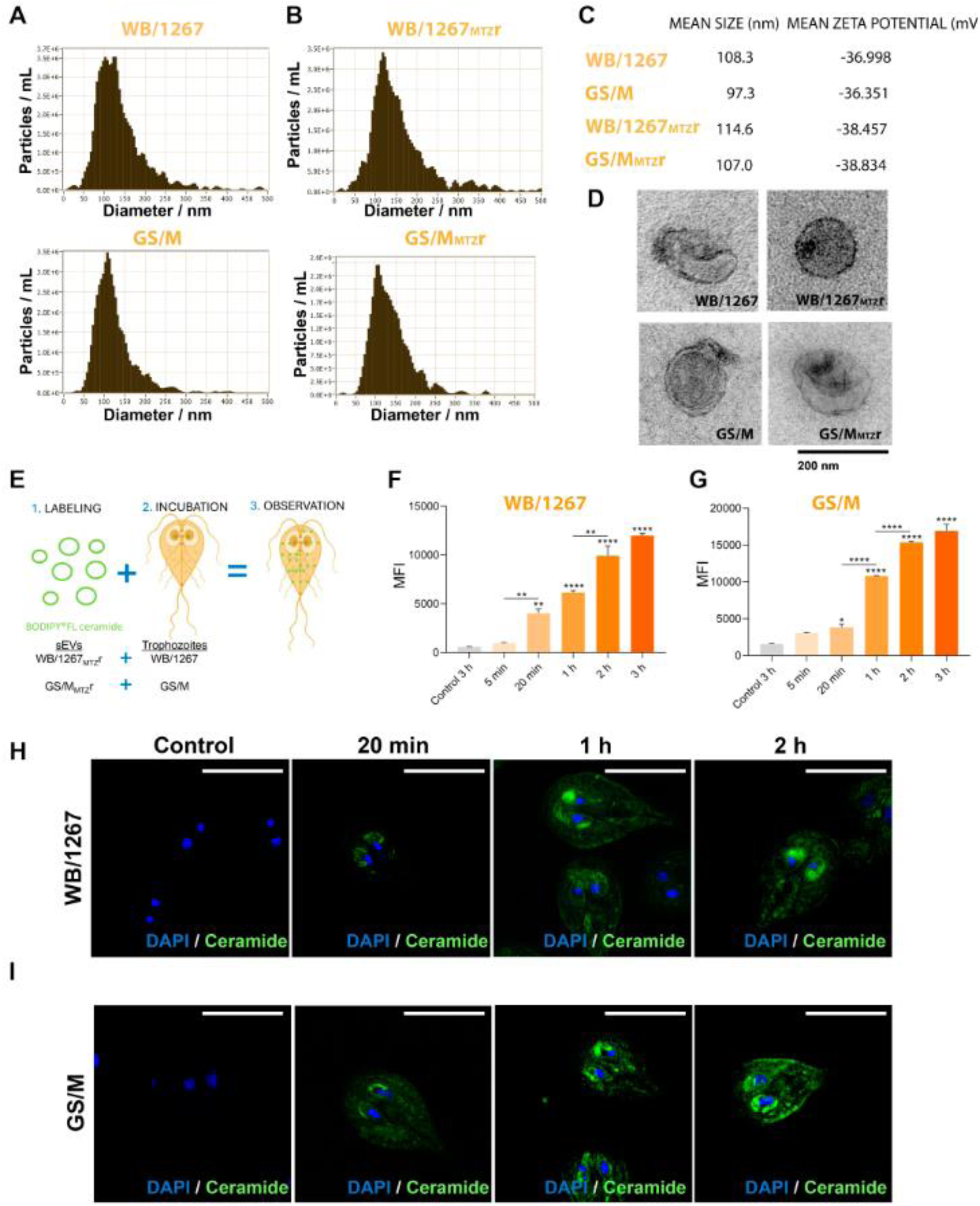
Characterization and functional analysis of small extracellular vesicles (sEVs) released by metronidazole-resistant (MtzR) *Giardia lamblia* clones. (A-B) Nanoparticle tracking analysis (NTA) of sEVs derived from wild type and MtzR clones WB/1267_MTZ_r and GS/M_MTZ_r. Mean particle sizes were around 100 nm for all samples, with uniform size distribution and no contaminants detected. (C) Zeta potential (ZP) measurements of sEVs showed consistent negative values across all samples, indicating electrostatic repulsion and colloidal stability. (D) Transmission electron microscopy (TEM) revealed characteristic cup-shaped structures with a mean diameter of approximately 100 nm. (E) Cartoon depicting the internalization of BODIPY FL-C5-ceramide-labeled sEVs from MtzR clones into wild type trophozoites. (F-G) Flow cytometry analyses of BODIPY-associated Median Fluorescence Intensity (MFI) disclose that RsEV uptake is time-dependent and reaches a saturation plateau after 2 hours. *p < 0.05; **p < 0.01, ***p < 0.001, ****p < 0.0001. (H-I) Super-resolution microscopy shows that ceramide (green) is localized to the endoplasmic reticulum (ER), perinuclear membranes (PNM), and peripheral vacuoles (PVs). Bars: 10 µm. Control experiments with disrupted sEVs or PBS showed no significant internalization.

To investigate whether *Giardia* RsEVs function as carriers between trophozoites, as previously demonstrated ^33^., RsEVs isolated from WB/1267_MTZ_r and GS/M_MTZ_r clones were labeled with green-fluorescent membrane probe BODIPY FL-C5-ceramide, a well-established marker for exosome membranes both *in vitro* and *in vivo* ^47^. Since *Giardia* cannot synthesize ceramide *de novo*, it must acquire this lipid from its environment ^48^. Thus, labeled RsEVs from WB/1267_MTZ_r were washed and incubated with WB/1267 wild type isolates at various times. The same procedure was performed using RsEVs from GS/M_MTZ_r cells and GS/M wild type isolates (Figure 2E). Flow cytometry analysis revealed a progressive increase in BODIPY-labeled ceramide uptake by *Giardia* during incubation. However, no significant changes were observed in RsEV internalization or ceramide incorporation between 2 and 3 hours of incubation (Figure 2F-G). Super-resolution microscopy observations showed that the ceramide contained within RsEV membranes was incorporated in a time-dependent manner, localizing in the perinuclear membranes (PNM), the endoplasmic reticulum (ER), and also detected at the point of internalization, specifically in the endo-lysosomal peripheral vacuoles (PVs) of WB/1267 and GS/M trophozoites up to the saturation point (2 h) (Figure 2H-I). All experiments’ controls included incubation with disrupted RsEVs from MtzR clones or PBS before labeling and incubation with WB/1267 or GS/M isolates. These findings demonstrate that RsEVs from MtzR *G. lamblia* clones can be internalized by wild type trophozoites, following a pathway like that of ceramide uptake ^49,50^.

### The internalization of RsEVs released by drug-resistant parasites influences the MTZ sensitivity of isogenic wild type cells

After confirming the transfer of RsEVs from MtzR trophozoites to isogenic wild type trophozoites, we assessed whether this incorporation affected MTZ resistance. Various RsEV-to-trophozoite ratios and incubation times were tested to optimize experimental conditions. As detailed in the STAR+METHODS section, the most substantial effect was observed with a 1000:1 RsEV-to-trophozoite ratio and four rounds of 2-hour incubations. Cellular uptake of RsEVs improved with increasing incubation times and was maximized through multiple incubation cycles with recovery periods between treatments (Figure 3A).

**Figure 3:**
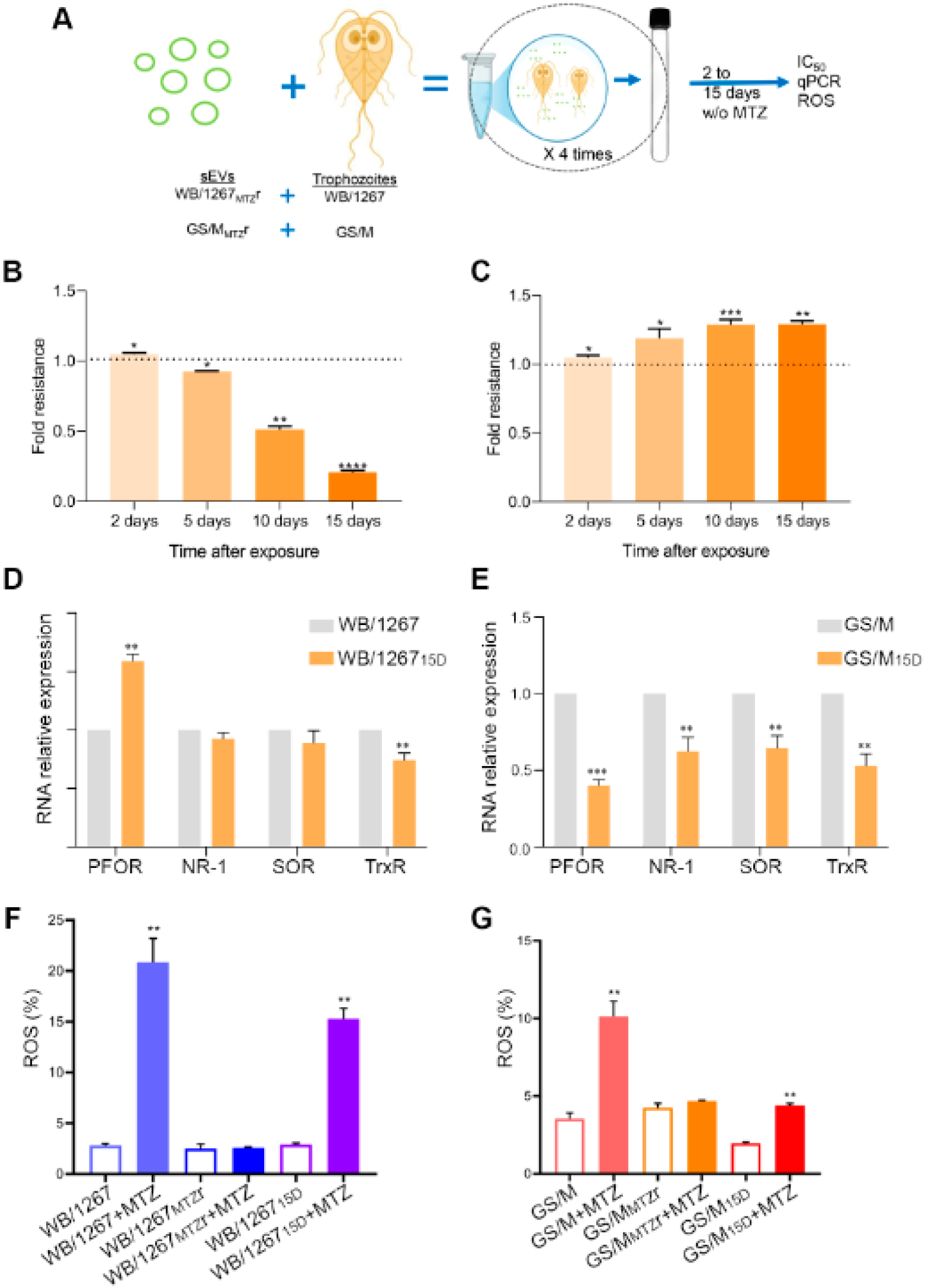
Effects of resistant small extracellular vesicles (RsEVs) on metronidazole (MTZ) sensitivity in isogenic wild type *Giardia lamblia* trophozoites. (A) Experimental optimization of RsEV incorporation using a 1000:1 RsEV-to-trophozoite ratio and four cycles of 2-hour incubations with recovery periods. (B-C) Phenotypic changes in MTZ sensitivity following RsEV uptake. WB/1267 wild type trophozoites treated with RsEVs from WB/1267_MTZ_r showed an 84% increase in MTZ sensitivity by day 15 (WB/1267_R15_), while GS/M trophozoites treated with RsEVs from GS/M_MTZ_r exhibited a 31% increase in MTZ resistance by the same time point (GS/M_R15_). (D-E) Differential expression of MTZ metabolism-related enzymes as assessed by qPCR. Pyruvate ferredoxin oxidoreductase (PFOR), nitroreductase-1 (NR-1), superoxide reductase (SOR), and thioredoxin reductase (TrxR). (F-G) Percentage of intracellular reactive oxygen species (ROS) production after treatment with 20 µM MTZ, compared to their wild type. *p < 0.05; **p < 0.01, ***p < 0.001, ****p < 0.0001.

Initially, we evaluated the impact of RsEVs from WB/1267_MTZ_r on the phenotype of WB/1267 wild type isolates. After 15 days of incubation without MTZ, the resulting cells, termed WB/1267_R15_, exhibited an 84% increase in MTZ sensitivity compared to the original WB/1267 isolates, despite an initial increase in MTZ resistance observed after two days of culture (Figure 3B).

In contrast, when RsEVs from GS/M_MTZ_r cells were incubated with GS/M wild type isolates under the same conditions, a significant increase in MTZ resistance was observed. By day 15, the GS/M_R15_ cells exhibited a 31% increase in MTZ resistance compared to the wild type GS/M isolates (Figure 3C). These contrasting responses between WB and GS genotypes suggest that the effect of sEV-mediated transfer of resistance traits highly depends on the genetic background of *G. lamblia* isolates.

### The differential expression of enzymes involved in MTZ metabolism might explain the variations in MTZ-resistance acquisition

To explore further the differences in MTZ resistance, we performed qPCR analysis to determine whether the increased MTZ sensitivity of WB/1267_R15_ and the increased MTZ resistance of GS/M_R15_ were associated with changes in mRNA expression of PFOR, NR-1, SOR, and TrxR. The results revealed higher PFOR expression and reduced TrxR expression in WB/1267_R15_ compared to the original WB/1267 isolate. The reduced expression of TrxR may decrease the cells’ ability to counteract MTZ-induced damage, contributing to their heightened sensitivity to the drug (Figure 3D). Conversely, GS/MR_15_ cells exhibited decreased expression of PFOR and NR-1, suggesting a reduced capacity to convert MTZ into its toxic intermediates—an established resistance mechanism. Furthermore, the reduced mRNA expression of SOR and TrxR implies that GS/M_R15_ trophozoites either require these enzymes less due to lower production of toxic intermediates or rely less on detoxification pathways involving these enzymes. Instead, they may utilize alternative mechanisms to manage oxidative stress and resist MTZ (Figure 3E). The differential expression patterns of these enzymes observed between WB/1267_R15_ and GS/M_R15_ provide a molecular basis for their opposed responses to MTZ. In WB/1267_R15_, increased sensitivity seemed linked to higher activation of MTZ and reduced detoxification capabilities. In contrast, GS/M_R15_ showed increased resistance through decreased drug activation and altered stress response mechanisms.

Since an increase in ROS production following MTZ treatment is ultimately responsible for trophozoite viability, we performed the H_2_DCFDA assay on WB/1267_MTZ_r, GS/M_MTZ_r, WB/1267_R15_, and GS/M_R15_ trophozoites, comparing their ROS production to that of the respective wild type WB/1267 or GS/M genotypes. The results showed that, although WB/1267_R15_ produced fewer ROS than WB/1267 after treatment with 20 µM MTZ, the level of ROS in WB/1267_R15_ was significantly higher than in the WB/1267_MTZ_r resistant clone, supporting the gaining of MTZ sensitivity in WB/1267_R15_ (Figure 3F). In contrast, GS/M_R15_ trophozoites showed significantly lower levels of ROS than GS/M, similar to the resistant GS/M_MTZ_r (Figure 3g). These findings support the acquired MTZ sensitivity in WB/1267_R15_, while GS/M_R15_ exhibits robust resistance, likely due to its reduced ROS generation, which is critical to MTZ’s toxicity. Overall, these results highlight the complexity of MTZ resistance in *G. lamblia* and suggest that resistance mechanisms can vary according to genotype-specific pathways.

### The variability on MTZ-resistance transmission might rely on the RsEV content and the genetic particularities of the recipient cell

The large genetic distances that separate *G. lamblia* genotypes suggest they represent distinct species, a conclusion reinforced by comparisons at the whole genome level of isolates representing Assemblages A, B, and E ^12,51,52^. The symptoms and the disease can vary depending on the genotype (genetic group) involved, with assemblages A and B causing similar gastrointestinal symptoms. However, studies have shown some differences in the clinical presentation, with genotype A being more asymptomatic and associated with longer-lasting, chronic infections, and genotype B has been often associated with more severe gastrointestinal issues and higher recurrence ^15,53–56^. In addition to the complexity of the diagnosis related to the symptoms, another challenge is the reported mixed infections, which ranged from 2.0% to 21.0% and were higher in less economically developed countries ^57–62^. Considering that MTZ resistance or sensitivity can be transmitted between trophozoites, we hypothesized that this information might also be exchanged between different genotypes. If so, the acquisition of MTZ sensitivity or resistance might be linked to the information carried by sEVs. To test the kinetic of the sEVs uptake, an enriched fraction of BODIPY FL-C5-ceramide-labeled RsEVs obtained from the WB/1267_MTZ_r clone was incubated with the wild type GS/M isolate while labeled RsEVs from GS/M_MTZ_r were incubated with WB/1267, following the protocol described above (Figure 4A). Super-resolution microscopy showed that RsEVs uptake increased over time in both scenarios (Figure 4B-C). However, by flow cytometry analysis, a significant reduction in uptake was observed when RsEVs from WB/1267_MTZ_r were incubated with GS/M trophozoites than the opposite combination (Figure 4B-C, graphic). These results become more evident when BODIPY-ceramide fluorescence is analyzed by flow cytometry, particularly when comparing genotypes (Figure S2).

**Figure 4.**
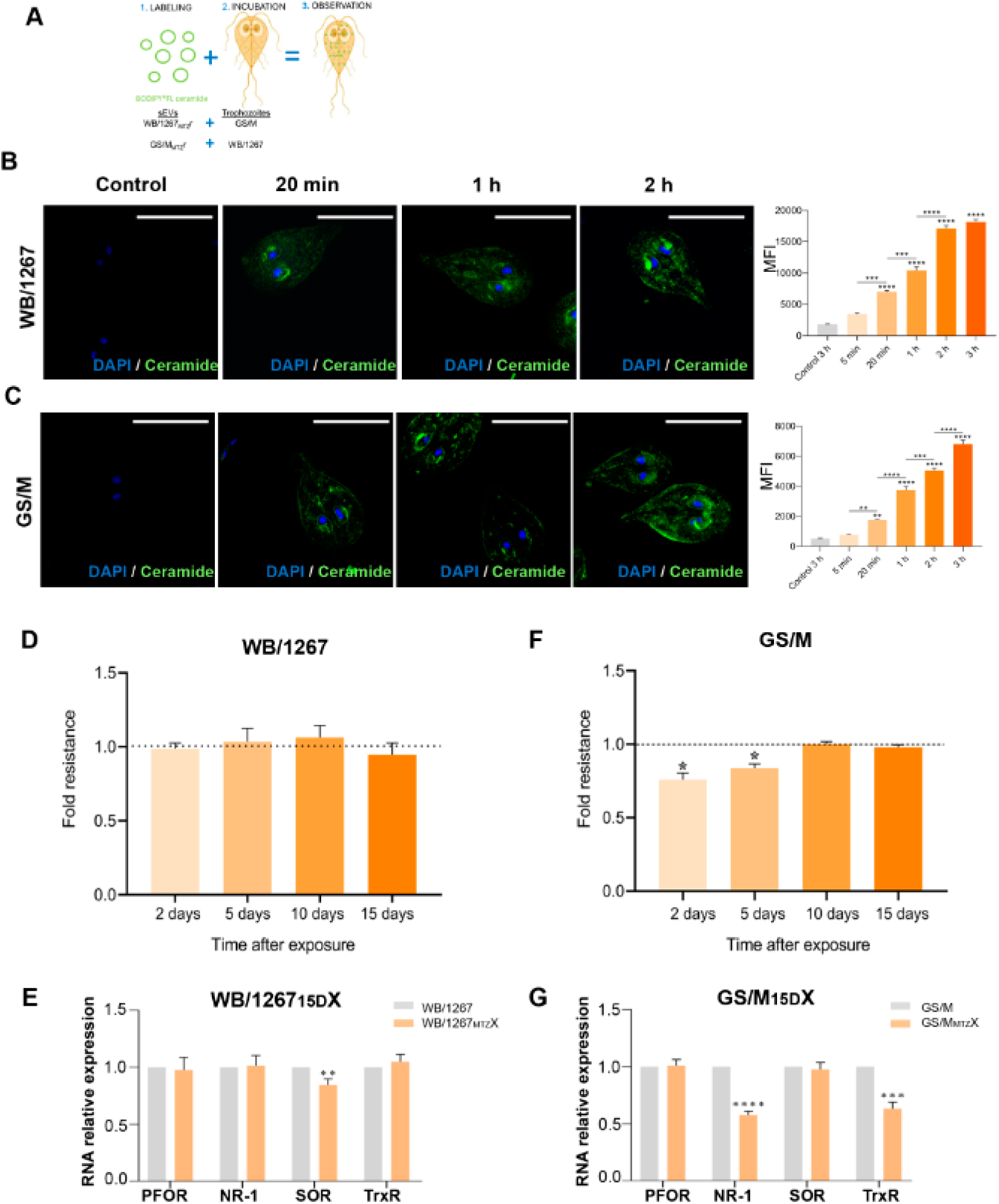
Analysis of the role of RsEVs in the transmission of metronidazole (MTZ) resistance between no isogenic *Giardia lamblia* genotypes. (A) Schematic representation of the experimental setup for RsEV uptake in which RsEVs from WB/1267_MTZ_r clone were incubated with the wild type GS/M isolate, while labeled RsEVs from GS/M_MTZ_r were incubated with WB/1267. (B-C) Super-resolution microscopy images show increased RsEV uptake over time in both experimental conditions. Bars: 10 µm. The graphics of flow cytometry analyses of BODIPY-associated Median Fluorescence Intensity (MFI) demonstrate a significant reduction in uptake when RsEVs from WB/1267_MTZ_r were incubated with GS/M trophozoites, compared to the reverse combination. (D) MTZ sensitivity was assessed by IC_50_ fold changes at different time points. No significant changes were observed when sEVs from GS/M_MTZ_r were incubated with WB/1267. (E) Enzyme expression in the derived WB/1267_R15_X strain shows no difference (except for SOR mRNA) compared to the wild type, suggesting the absence of effective resistance adaptations. (F) Increased sensitivity to MTZ (lower IC_50_) in GS/M trophozoites at early time points when incubated with sEVs from WB/1267_MTZ_r. (G) Reduced expression of NR-1 and TrxR in the GS/M_R15_X strain, indicate partial or incomplete resistance mechanisms. Pyruvate ferredoxin oxidoreductase (PFOR), nitroreductase-1 (NR-1), superoxide reductase (SOR), and thioredoxin reductase (TrxR). *p < 0.05; **p < 0.01, ***p < 0.001, ****p < 0.0001.

When the effect of MTZ was assessed using the protocol settled and the fold changes in the IC_50_ at different time points were measured, no significant changes were seen when RsEVs from GS/M_MTZ_r were incubated with WB/1267 (Figure 4D). In this sense, the derived WB/1267_R15_X strain demonstrated no difference in sensitivity or enzyme expression compared to the wild type, indicating that resistance adaptations are absent or ineffective (Figure 4D). However, an increased sensitivity to MTZ (lower IC_50_) was observed in GS/M trophozoites incubated with RsEVs from WB/1267_MTZ_r only at early time points (Figure 4F) and the GS/M_R15_X strain showed only reduced expression of NR-1 and TrxR (Figure 4G), implying partial or incomplete resistance mechanisms. These results suggest that the information carried by RsEVs plays a critical role in transferring sEV-mediated MTZ resistance, potentially influencing the recipient cells’ response to the drug. However, the genotype strongly influences this process, underscoring the intricate and genotype-specific nature of drug resistance mechanisms in *G. lamblia*. These findings highlight the potential importance of RsEVs as mediators of resistance and the limitations of inter-genotype resistance transmission. Further investigation is essential to unravel the specific mechanisms by which RsEVs contribute to resistance transfer and their interplay with the genetic background of the parasite.

## Discussion

This study showed, for the first time, that the small extracellular vesicles in *G. lamblia* can transmit information between parasites that control their response to MTZ. Equally important, it proved that this information can be overwhelmed by features inherent to the genotype.

One approach to studying drug resistance involves using isolated strains from patients who experienced treatment failure with MTZ. However, it is challenging to distinguish between reinfection and actual drug resistance in infected patients. Moreover, isolates are complex to culture, and growth rates may differ between isolates and assemblages ^63^. Consequently, the preferred method is the *in vitro* induction of resistance by subculturing parasites under increasing sublethal concentrations of MTZ. This approach mimics real-world scenarios, as one cause of drug resistance is the discontinuation of treatment or suboptimal drug dose in the treatment protocol ^64^. Additionally, *in vitro*-generated resistance clones allow for controlled experimentation and growth analysis.

This study provides the first evidence of laboratory-induced MTZ resistance in *G. lamblia* trophozoites of genotype B, specifically in the GS/M_MTZ_r clone. Starting with similar MTZ sensitivity levels as genotype A, the GS/M_MTZ_r clone developed 2.5 times greater resistance. This heightened resistance aligns with previous findings that genotype B exhibits distinct capabilities in managing oxidative and oxidative stress ^65^.

Our recent findings demonstrated that *Giardia* sEVs transfer sRNAs, including shared and distinct biotypes, between trophozoites of the same genotype ^33^, highlighting the potential role of sEVs as carriers of specific molecules with distinct biological information. Based on previous studies linking exosomes (now referred to as small extracellular vesicles, or sEVs) to drug resistance in tumor cells ^66–69^, we hypothesized that sEVs could specifically mediate the transmission of MTZ resistance in *G. lamblia*. Recent metagenomic studies have identified antibiotic-resistance genes within bacterial EVs, supporting these findings ^70^. Additionally, EVs derived from drug-resistant *Leishmania* parasites have been shown to transfer episomal DNA containing drug-resistance genes ^29^. This transfer enables recipient parasites to exhibit enhanced growth and improved oxidative stress management. The authors of these *Leishmania* studies concluded that parasites exploit EVs—predominantly those under the 200 nm size threshold— to propagate drug-resistance genes as part of episomal amplification ^29^. This constitutes a clear demonstration of EV-mediated horizontal gene transfer in eukaryotic parasites and represents an alternative mechanism for drug resistance in eukaryotic cells.

Here, we demonstrated that RsEVs released by drug-resistant parasites can efficiently alter the drug-sensitivity phenotype of recipient parasites after four days of exposure, revealing a rapid adaptation and modulation of MTZ metabolism. Remarkably, the recipient trophozoites, WB/1267_15D_ and GS/M_15D_, exhibited divergent responses to RsEV-mediated molecular exchange: WB/1267_15D_ lost resistance to MTZ, while GS/M_15D_ displayed enhanced resistance. The MTZ sensitivity observed in WB/1267_15D_ trophozoites seemed linked to higher MTZ metabolism and reduced detoxification capacity. In contrast, GS/M_15D_ trophozoites displayed enhanced resistance due to decreased drug activation and altered oxidative stress response pathways. These results suggest that RsEV-mediated molecular exchange induces rapid and genotype-specific adaptations in *G. lamblia*, with divergent effects on drug sensitivity determined by the RsEVs information or the recipient’s metabolic and stress response pathways.

In our investigation of RsEVs’ role in redox control and metabolic enzyme regulation related to MTZ, we found that the GS/M trophozoites incubated with RsEVs from WB/1267MTZr demonstrated heightened sensitivity to MTZ at early stages, though this effect diminished after 15 days. Furthermore, the incubation of RsEVs from GS/M_MTZ_r with the wild type WB/1267 isolate did not produce significant changes. These findings indicate that the transfer of resistance through RsEVs is genotype-dependent and not universally effective, highlighting the complexity of resistance mechanisms in these parasites.

The observed differences in resistance between GS/M and WB/1267 strains to MTZ after incubation with RsEVs from isogenic clones suggest that GS/M strains possess a combination of genetic, metabolic, and antioxidant factors that enhance their ability to resist the drug. These factors could include genetic mutations, higher levels of MTZ detoxifying enzymes, and efficient antioxidant systems. In contrast, the increased sensitivity of WB/1267_R15_ to MTZ following exposure to RsEVs from WB/1267_MTZ_r suggests that the integration of RsEV-derived cargo may disrupt the redox balance, alter enzyme expression, and increase of ROS production, ultimately making the cells more vulnerable to MTZ. These findings highlight the complex, genotype-specific mechanisms underlying MTZ resistance and the impact of RsEV-mediated molecular exchange on drug sensitivity. Further investigation is needed to better understand the interplay between genetic background and sEV-mediated resistance transfer.

Resistance to MTZ has been observed in *G. lamblia* isolates from patients and *in vitro*-derived clones of genotype A. Previous studies have underscored the importance of controlling MTZ metabolism and managing the cellular response to toxic molecules, such as reactive oxygen species. However, the mechanisms underlying the acquisition of this resistance remain largely unexplored. Although poorly understood in eukaryotic parasites, epigenetic mechanisms may provide a plausible explanation for the rapid shifts in MTZ sensitivity observed when wild type trophozoites are exposed to RsEVs from isogenic or non-isogenic clones. Emerging evidence suggests that functional crosstalk between the modification of histones and chromatin remodeling is essential in transcriptional regulation and cellular decision-making. Our preliminary findings indicate that histone 3 modifications significantly regulate MTZ resistance in the *G. lamblia* clones studied, with distinct differences between genotypes and resistant clones (Luna Pizarro et al., results not shown). Additionally, sRNAs carried by *G. lamblia* sEVs may act as transcriptional regulators in recipient cells, as demonstrated for tsRNAs from *Trichomonas* vaginalis ^71^. Further studies employing approaches such as ChIP-Seq and RNA-Seq will be essential to provide conclusive insights into these mechanisms.

Since many pathogens share conserved epigenetic pathways, targeting these pathways may reveal novel drug targets, potentially leading to broad-spectrum antiparasitic agents. The emergence of drug-resistant microorganisms represents a growing threat to global health, particularly among immunocompromised individuals. Despite advances in prevention and diagnostics, MTZ resistance in *G. lamblia* continues to cause significant morbidity and mortality. Current therapeutic options are limited to a single primary drug, MTZ, emphasizing the urgent need for alternative treatments. The availability of MTZ-resistant *G. lamblia* clones presents an opportunity to screen already approved drugs, accelerating the discovery of new therapies to combat drug-resistant giardiasis and other protozoan infections.

Addressing MTZ resistance in *G. lamblia* requires a multidisciplinary approach integrating molecular, epigenetic, and therapeutic studies. Unraveling the mechanisms of resistance acquisition and transmission advances our understanding of giardiasis. It informs the broader fight against drug-resistant pathogens, underscoring the critical need for innovation in antiparasitic drug development.

### Limitations of the study

While the study demonstrates genotype-specific adaptations in response to MTZ, it also showed that resistance mechanisms are not universally effective across all genotypes. This underlines the genotype-dependent nature of MTZ resistance transfer through sEVs, indicating the need for further investigation to fully understand the genetic background’s role in resistance transmission. The findings also suggest that specific pathways, such as ROS production and metabolic enzyme regulation, are involved, but these mechanisms require further characterization. The study acknowledges the potential role of epigenetic mechanisms, including histone modifications, in regulating MTZ resistance. Although preliminary findings point to the significance of these mechanisms, the detailed molecular pathways remain unexplored. Further research employing techniques such as ChIP-Seq, proteomics, and RNA-Seq is essential to elucidate the role of epigenetics in MTZ resistance. Finally, the interaction between *G. lamblia* and the host immune system or gut environment, which may also influence resistance mechanisms, was not addressed in this study and warrants further exploration.

## RESOURCE AVAILABILITY

### Lead contact

Further information and requests for resources and reagents should be directed to and will be fulfilled by the lead contact, María Carolina Touz.

### Materials availability

This study did not generate new unique reagents.

## EXPERIMENTAL MODEL AND SUBJECT DETAILS

### Cell lines and cell culture

Axenic cultures of *Giardia* trophozoites of isolates WB clone 1267 (ATCC 50582, Assemblage A) and GS/M (ATCC 50581, Assemblage BIV) were purchased at American Type Culture Collection (www.atcc.org, accessed on 1 March 2008). Trophozoites were routinely grown in 16 mL screw-cap tubes (NuncTM, ThermoFisher Scientific, Waltham, MA, USA) in TYI-S33 medium, supplemented with 10% adult bovine serum (Euroclone, Pero, Italy) and bovine bile (Sigma-Aldrich S.R.L., Milan, Italy) ^72^ at 37 °C. Log-phase cultures were harvested after cooling the culture vials on ice for 15 min and centrifugation at 700 × g for 10 min.

### Production of metronidazole-resistant *Giardia* clones

Metronidazole (MTZ)-resistant *Giardia* clones were generated by gradually exposing sensitive trophozoites to increasing sublethal concentrations of MTZ, followed by selection and subcloning. Briefly, *Giardia* trophozoites from strains WB 1267 (ATCC 50582, Assemblage A) and GS/M (ATCC 50581, Assemblage BIV) were cultured in complete growth medium supplemented with sublethal concentrations of MTZ (Cat. No. M3761, Sigma-Aldrich). The adaptation process involved a stepwise protocol, beginning with an initial sublethal concentration of 0.25 μM and progressing through successive rounds of culture in higher concentrations up to 6.4 μM and 10.5 μM for strains WB/1267_MTZ_r and GS/M_MTZ_r, respectively. At each stage, adapted trophozoites were selected and continuously cultured under these conditions for 730 days, with regular monitoring to assess cell viability.

## METHOD DETAILS

### Cell viability assay

To assess the cytotoxic effect of MTZ, the 3-(4,5-dimethylthiazol-2-yl)-2,5-diphenyltetrazolium bromide (MTT) colorimetric assay was performed as previously described ^73^. Briefly, trophozoites from WB/1267, GS/M, WB/1267_MTZ_r, GS/M_MTZ_r, WB/1267_R15_, and GS/M_R15_ isolates were seeded at a density of 5 × 10⁵ cells per well in 150 μL of complete growth medium in 96-well plates. An additional 150 μL of medium containing serial dilutions of MTZ, previously dissolved in DMSO (final concentration 0.5% v/v, as this concentration showed no adverse effects on cell growth), was added. Following anaerobic incubation at 37°C for 48 hours, the plates were centrifuged at 500 x g for 10 min. Subsequently, the cells were washed three times with 1 x PBS by centrifugation, and 20 μL of a 5 mg/mL MTT solution in sterile PBS was added to each well, followed by further incubation for 4 hours at 37°C. After the removal of the supernatants, 100 μL of DMSO was added to solubilize the purple formazan crystals produced by metabolically viable cells. Absorbance was measured at 570 nm using a Model 680 microplate reader (Bio-Rad, USA). Cytotoxicity percentages were calculated relative to DMSO-treated control cells, which were considered 100% viable. The percentage of cytotoxic activity was determined using the following formula: cytotoxicity (%) = [1 − (OD of treated cells − OD of DMSO) / (OD of control cells − OD of DMSO)] × 100, where OD refers to optical density. Half-maximal inhibitory concentrations (IC_50_), defined as the concentrations required to inhibit 50% of cell proliferation, were determined from the mean values obtained from replicate wells across three independent experiments.

The resistance fold was calculated by comparing the IC_50_ values of sensitive and resistant trophozoites using the following formula: Resistance Fold (RF) = IC_50_ of resistant cells / IC_50_ of sensitive cells. RF values greater than 1 indicate increased tolerance to MTZ, values less than 1 indicate reduced tolerance, and values equal to 1 suggest no change in tolerance.

### RT-qPCR analysis

The expression of key genes involved in MTZ metabolism and reactive oxygen species detoxification was assessed using RT-qPCR. Briefly, trophozoites from the WB/1267, GS/M, WB/1267_MTZ_r, GS/M_MTZ_r, WB/1267_R15_, and GS/M_R15_ isolates were homogenized in Trizol reagent (Cat. No. 15596026, Thermo Fisher Scientific Inc.). Total RNA was extracted using the SV Total RNA Isolation System (Cat. No. Z3100, Promega, Madison, WI, USA) following the manufacturer’s protocol. Two micrograms of total RNA were reverse-transcribed using M-MLV Reverse Transcriptase (Cat. No. A3802, Promega). The cDNA was analyzed for the PFOR-2, NR-1, SOR, and TrxR genes via real-time PCR using SYBR Green Master Mix (Cat. No. QR100-1KT, Thermo Fisher Scientific Inc.), with 100 ng of total RNA equivalent as single-stranded cDNA and 800 nM of each amplification primer in a 20 μL reaction volume. Specific primers for each gene were designed using Primer Express software (Applied Biosystems, Foster City, CA, USA): PFOR-2 Fw (CATGAACACGGAGCAGAGGT) and PFOR-2 Rv (GAGCCCCTGAAGAACCTTCC); NR-1 Fw (CGAGACAAAGGTAGTGGCGT) and NR-1 Rv (CTGCCGGTGGATCTGTCTTT); SOR Fw (GAGGACCAAGGAGAAGCACG) and SOR Rv (TTGCCCTCCTTAGTGATGCC); TrxR Fw (CTCGCTGACGCCCTTATCAT) and TrxR Rv (GACACCCCCTTTTGCCAGTA). Runs were carried out on a standard 7500 system (Applied Biosystems) using a QuantStudio3 device (Applied Biosystems, Foster City, CA, USA) and QuantStudio™ Design & Analysis Software v1.5.2. The RT-qPCR conditions were as follows: 50°C for 2 min, 95°C for 10 min, followed by 40 cycles of 95°C for 15 s and 60°C for 1 min. Gene expression was normalized to the housekeeping gene 18S (Fw: AAGACCGCCTCTGTCAATAA, Rv: GTTTACGGCCGGGAATACG) and calculated using the comparative ΔΔCt method. Melting curve analysis was performed to confirm the specificity of the qPCR products. These assays were conducted in triplicate with duplicate reactions. All DNA oligonucleotides were purchased from Macrogen (Macrogen, Seoul, Republic of Korea).

### Measurement of Reactive Oxygen Species (ROS)

Following the manufacturer’s instructions, the Image-iT LIVE Green Reactive Oxygen Species Detection Kit (Invitrogen, Massachusetts, USA) was used to assess intracellular ROS generation. Flow cytometry (FACS Canto II, Becton & Dickinson, New Jersey, USA) was employed to quantify ROS levels using the fluorescent probe 2’,7’-dichlorodihydrofluorescein diacetate (H_2_DCFDA), which is oxidized to 2’,7’-dichlorofluorescein (DCF) in the presence of ROS, emitting fluorescence that is proportional to the oxidative capacity of reactive species. ROS levels were measured in the following isolates: WB/1267, GS/M, WB/1267_MTZ_r, GS/M_MTZ_r, WB/1267_R15_, and GS/M_R15_, after a 48-hour treatment with 20 µM metronidazole or without the drug as a control.

### Isolation and purification of small extracellular vesicles (sEVs)

Enriched sEVs were obtained using differential ultracentrifugation from the supernatant of trophozoites, as we described ^34^. Briefly, 7 × 10^7^ trophozoites recovered from the monolayer were washed twice with warm PBS 1X (37 °C). To avoid exosomal contamination from other sources, the trophozoites were incubated in TYI-S-33 medium without serum and bovine bile (TYI-S-33/-sbb) for 4 h at 37 °C before isolation. Then, the parasites were removed by centrifugation at 1455× g for 15 min and the supernatant recovered. After centrifugation, the supernatant was filtered through a 0.11-µm filter (Millipore) to discard high-size vesicles. To obtain the sEVs fraction, the filtered elution was subsequently pelleted at 100,000× g for 180 min using a 60Ti rotor (Beckman-coulter L-70 Ultracentrifuge). The pellet was then washed with PBS and pelleted again at 100,000× g in the same ultracentrifuge.

### Identification and characterization of small extracellular vesicles (sEVs)

Nanoparticle tracking analysis (NTA) was used to determine the sEVs size distribution and concentration with a ZetaView PMX-230 Twin Laser (Particle Metrix, Germany) device according to the MISEV 2023 guidelines ^74^. Briefly, sEV-enriched samples were re-suspended in 0.11-μm-filtered PBS 1X and diluted to achieve a particle concentration within the detection range (20– 100 particles/frame) before measurement. Samples were manually injected into the instrument using a 1-mL syringe. The measurements were taken at 11 different positions, with video quality set to medium and camera sensitivity set to 80. Data analysis was performed using ZetaView® software (version 8.05.16 SP7), with a minimum particle size of 10, a maximum size of 1000, and a minimum brightness of 30. All measurements were conducted at room temperature (25°C). A JEOL 1230 transmission electron was used for the visualization of exosomes. For negative staining electron microscopy, sEVs were diluted in PBS 1X, applied to copper grids, and incubated for 15 minutes at room temperature. Excess liquid was removed by blotting. The grids were then stained with 2% uranyl acetate (w/v) (Merck, Darmstadt, Germany) for 30 seconds and observed as previously described ^34^.

### Small extracellular vesicles internalization assays

The uptake of sEVs by trophozoites was visualized using super-resolution microscopy and assessed by flow cytometry after labeling the sEVs with BODIPY FL-C5-ceramide (Cat. No. D3521, Invitrogen™), following the manufacturer’s instructions, with the exception that excess dye was removed by ultracentrifugation. Briefly, enriched sEVs were suspended in 100 µL of PBS per labeling reaction, and 1 µL of a 1 mM dye stock solution was added to the samples. After mixing, the samples were incubated at 37°C for 30 minutes, protected from light. Unincorporated dye was removed by ultracentrifugation at 100,000× g for 180 minutes using a 60Ti rotor (Beckman Coulter L-70 Ultracentrifuge). The pellet containing the freshly labeled sEVs was recovered and used in uptake assays. As a control for non-specific staining, the same procedure was applied to autoclaved sEVs.

For uptake assays, trophozoites were harvested by chilling culture tubes on ice, washed three times in PBS (pH 7.4), counted using a hemocytometer, and adjusted to the required concentration. 1 × 10^5 trophozoites were incubated with BODIPY FL-C5-ceramide-labeled sEVs for 20 minutes, 1 hour, or 2 hours at 37°C. A control incubation was performed with trophozoites and autoclaved labeled sEVs for 2 hours at 37°C. After incubation, trophozoites were centrifuged at 700× g for 15 minutes at 4°C. The supernatant was discarded, and the pellet was resuspended in PBS 1X. Trophozoites were placed onto Poly-L-Lysine-coated immunofluorescence slides and incubated at 37°C for 1 hour to allow cell adherence. The slides were then washed with 1× PBS, fixed with 4% paraformaldehyde for 20 minutes at room temperature, and washed twice. The trophozoites were then incubated with DAPI and mounted using FluoSafe® (Sigma). Fluorescence was visualized using a super-resolution confocal microscope ZEISS LSM 980 with Airyscan 2. Images were processed using Fiji software ^75^. For flow cytometry analysis, trophozoites were incubated with BODIPY FL-C5-ceramide-labeled sEVs for 5 minutes, 20 minutes, 1 hour, 2 hours, or 3 hours at 37°C. A control incubation with trophozoites and autoclaved labeled sEVs was carried out for 3 hours at 37°C. After incubation, trophozoites were centrifuged at 700× g for 15 minutes at 4°C, the supernatant discarded, and the pellet resuspended in PBS 1X. The mean fluorescence intensity (MFI) was measured using a BD FACSCanto™ flow cytometer (BD Biosciences), and the relative distribution of 10,000 cells was analyzed using FlowJo software (version 7.6.2, Tree Star, Inc., OR, USA).“

### Production of metronidazole-resistant *Giardia* clones by small extracellular vesicles (sEVs)

Changes in the MTZ resistance fold induced by the uptake of RsEVs in sensible trophozoites were assessed after successive rounds of exposure. The concentration of RsEVs was determined using nanoparticle tracking analysis, and trophozoites were harvested and counted as described previously. Briefly, 1 × 10¹⁰ RsEVs were incubated with 1 × 10⁷ sensitive trophozoites (cells/RsEVs ratio: 1/1000) for 2 hours at 37°C in a final volume of 2 mL (PBS 1× was added as needed). After the exposure period, the contents of the tube were transferred to a 16 mL tube, supplemented with a complete growth medium, and incubated at 37°C for 24 hours to allow recovery of the treated cells. This procedure was repeated for a total of four rounds. Following the four rounds of treatment, the cells were allowed to recover for 2, 5, 10, or 15 days. The MTZ IC_50_ of the treated trophozoites was determined at each recovery time point, and the corresponding resistance fold was calculated by comparing the IC_50_ of trophozoites treated with RsEVs to that of cells treated with sEVs.

### Statistical analyses

All statistical analyses were performed using GraphPad Prism 9 (GraphPad Software Inc., USA, www.graphpad.com). Data are presented as the mean ± standard error of the mean (SEM) from three independent experiments. Statistical significance was assessed using paired and unpaired one-tailed Student’s t-tests, with a significance threshold set at p ≤ 0.05. Additionally, for the small extracellular vesicles (sEVs) internalization assays, a one-way analysis of variance (ANOVA) was conducted to compare internalization levels across experimental groups. Post hoc comparisons were performed using Tukey’s multiple comparisons test, with a significance level of p ≤ 0.05.

## Declaration of generative AI and AI-assisted technologies in the writing process

During the preparation of this work, the author(s) utilized ChatGPT (OpenAI, 2024) and *Grammarly* (https://www.grammarly.com) to assist with English language refinement and readability. The author(s) carefully reviewed and edited all content as necessary and assume(s) full responsibility for the final version of the publication. Schemes for the figures were created using BioRender.com

## DECLARATION OF INTERESTS

The authors declare no competing interests.

## Acknowledgments

The authors thank Andrea Vanina Pellegrini, Laura Montroull, María Silvina Ferrer, and Joaquín José Nigro for technical assistance. We also thank Dr. Gonzalo Quassollo and Dr. Carolina Leimgruber for their assistance in confocal and electron microscopy, respectively We thank GAVE (Grupo Argentino de Vesículas Extracelulares, https://sites.google.com/view/gavear/p%C3%A1gina-principal) for their valuable suggestions and insights, which significantly enriched the discussion of this study. Dr. María Eugenia Santana and Marcela Cucher (Institute of Research on Microbiology and Medical Parasitology (IMPAM), School of Medicine, University of Buenos Aires) help with the NTA. We acknowledge the use of ChatGPT (OpenAI, 2024) and Grammarly for assistance in English language editing and proofreading of this manuscript. This work was supported by the National Agency for the Promotion of Science and Technology, grant numbers PICT-2021-I-A-00056 (M.C.T.), PICT Aplicado tipo II 2021-73 (M.C.T.), PICT 2019-0154 (A.S.R), and the Florencio Fiorini Foundation Grant 2024 and Universidad Católica de Córdoba (J.L.). All the authors were supported either by the National Agency for the Promotion of Science and Technology or the National Council on Scientific and Technical Research (CONICET), Argentina.

## Author contributions

Conceptualization, M.C.T. and G.L.P.; methodology, G.L.P., J.L., N.S., R.P, L.D.P, and C.C.; investigation, M.C.T., G.L.P., J.L., C.F., and A.S.R.; formal analysis, M.C.T., G.L.P, J.L.; writing – original draft, M.C.T.; writing – review & editing, M.C.T. G.L.P., J.L., C.F., and A.S.R.; supervision, C.F. and M.C.T.; funding acquisition, M.C.T., J.L., and A.S.R

## Notes

### Competing Interest Statement

The authors have declared no competing interest.

