## Supplementary figures 1 and 2 for "Genotype-specific roles of small extracellular vesicles in modulating metronidazole resistance in *Giardia lamblia*"

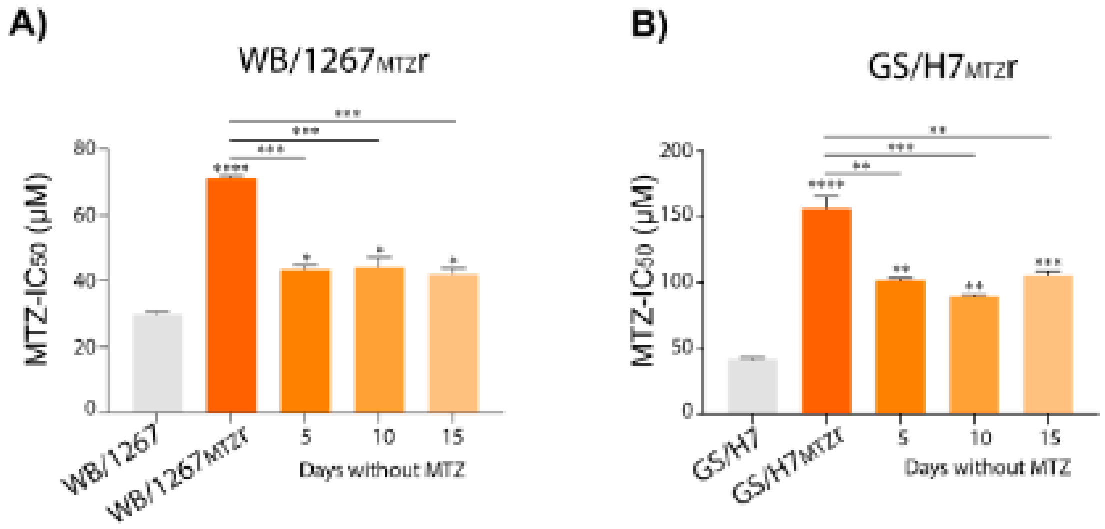


**Supplementary Figure 1**: **Assessment of metronidazole (MTZ) resistance stability in *Giardia lamblia* clones WB/1267$_{MTZ}$r and GS/M$_{MTZ}$r during culture without drug selection.** MTZ-resistant clones were cultured for 5, 10, and 15 days without MTZ to evaluate the potential for reversion to drug sensitivity. $IC_{50}$ values for MTZ were calculated at each time point. Significance was determined by comparing the clones cultured without the drug and the wild-type cells (* over each bar) and between the MTZr cells and the drug-free culture lines (* over the lines). Statistical significance levels are: *p < 0.05; **p < 0.01, ***p < 0.001.

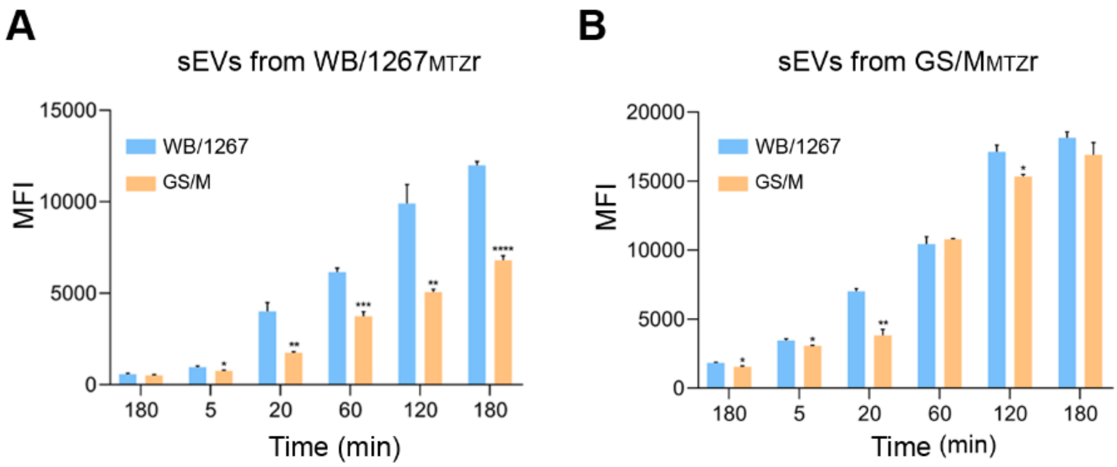


**Supplementary Figure 2**: **Analysis of variations in RsEV uptake by genotype.** (A-B) Flow cytometry analysis of BODIPY-associated Median Fluorescence Intensity (MFI) reveals a significant reduction in RsEV uptake when vesicles derived from WB/1267MTZr trophozoites are incubated with GS/M trophozoites, compared to the reverse combination. Statistical analysis was performed using one-way analysis of variance (ANOVA). *p < 0.05; **p < 0.01, ***p < 0.001, ****p < 0.0001.
